# Sulfonamide resistance gene *sul4* is hosted by common wastewater sludge bacteria and found in various newly described contexts and hosts including clinically relevant species

**DOI:** 10.1101/2025.01.22.634252

**Authors:** Melina Markkanen, Denise Pezzutto, Marko Virta, Antti Karkman

## Abstract

The introduction of the first broad-spectrum antibiotics, sulfonamide drugs fundamentally revolutionized medicine in the 1930s. Shortly after and ever since sulfonamide resistance genes (*sul* genes) have been widely detected. Still, the most recent variant of these genes *sul4*, was first described only in 2017 and its host range and transmission mechanisms are still largely unknown. Here we applied PacBio long-read metagenomic sequencing and bacterial methylation signals to investigate the genetic contexts and bacterial carriage of the *sul4* gene in wastewater. Furthermore, we extended our description of *sul4* carriers to previously published data sets. Our results indicate that *sul4* is prominently found in sludge and hosted by various bacteria such as the species from the phyla Myxococcota and Chloroflexota and genera *Trichlorobacter*, and *Desulfobacillus*, which are commonly found in activated sludge. Additionally, according to our results, *sul4* has already spread into multiple strains of opportunistic human pathogens, such as *Aeromonas* and *Moraxella* in addition to the previously described *Salmonella*. The sequence region flanking *sul4* included a truncated *fol*K gene, and an ISCR28-element, and exhibited a high degree of conservation across the investigated sequences. Furthermore, the module was associated with various integron integrase genes. Also, other mobility-related elements that could further increase the likelihood of *sul4* mobilization were detected. Altogether, our results describing the *sul4* hosts of bacteria from distant lineages indicate the efficient mobility of *sul4* by genetic elements that traverse both clinical and environmental bacteria. Finally, we suggest that wastewater may provide favorable conditions for such horizontal gene transfer events.

**Importance:** Antibiotic resistance is an ancient phenomenon and a common trait for many environmental bacteria. However, human activities in the post-antibiotic era, coupled with the bacteria’s ability to exchange genetic material across different lineages, have drastically increased the spread of resistance traits among bacteria from various niches. The primary concern is the resistance genes encoded by infections causing pathogens, already causing over one million deaths annually and indirectly contributing to nearly four million more. Therefore, understanding the bacteria that harbor ARGs and the genetic mechanisms driving their mobilization is crucial for understanding the dynamics and emerging trends of resistance. Here, we focus on revealing these crucial aspects of the newly discovered sulfonamide resistance gene *sul4*. Given the limitations of the metagenomic approach in linking the functional genes to their host genomes, the significance of our research lies in our workflow that allows this linkage by identification of shared methylation profiles.

## Introduction

Increased antibiotic resistance threatens our current dependence on effective antibiotics in medicine, animal husbandry, and food production systems (Mulchandani et al., 2023; Murray et al., 2022). Besides the accumulation of point mutations in bacterial DNA, resistance may emerge by horizontal transfer of antibiotic resistance genes (ARGs) mediated by mobile genetic elements (Larsson & Flach, 2022) such as plasmids (Nielsen et al., 2022; Rozwandowicz et al., 2018), IS-elements (He et al., 2019) and resistance gene cassettes encoded by integrons (Loot et al., 2024). Integrons are genetic elements that capture and express genes as cassettes obtained via site-specific recombination at *attI* and *attC* sites and mediated by an integrase gene (Loot et al., 2024). ISCR (insertion sequence common regions) are recently discovered and expanding group of IS91-like transposable elements that can translocate adjacent genes such as ARGs via a rolling circle mechanism by *ori*IS (origin of replication) and a *ter*IS (replication terminator) terminal sequences (Toleman et al., 2006; Yuan et al., 2024). Prophage-mediated spread of ARGs has also been recognized, though its importance requires further investigation (Calero-Cáceres et al., 2019; Wang et al., 2024). The transfer of ARGs carried by environmental bacteria into clinically significant bacterial hosts has been accelerated by human activities during the past decades (Collignon et al., 2018; Finley et al., 2013). Therefore, besides monitoring the occurrence of known resistance genes and identifying newly emerging ARGs, it is crucially important to study their surrounding genetic context and the bacteria that carry them. This aids in assessing their potential as high-risk ARGs that could migrate to new hosts and contexts (Larsson & Flach, 2022; Martínez et al., 2014).

In 2017, Razavi and colleagues made the first discovery of the novel gene *sul4*, which confers resistance to sulfonamide antibiotics (Razavi et al., 2017). This finding came decades after the identification of earlier *sul* gene variants: *sul1* in 1975 (Wise & Abou Donia, 1975), *sul2* in the year 1980 (Radstrom & Swedberg, 1988), and *sul3* in the year 2003 (Perreten & Boerlin, 2003). Ever since the introduction of sulfonamide antibiotics in 1935, they have been widely used in human medicine (Ovung & Bhattacharyya, 2019). Moreover, they are among the five most used antibiotics in veterinary medicine for treating livestock diseases, such as gastrointestinal and respiratory infections, and in some cases, as growth promoters (Mulchandani et al., 2023) and specifically more used in many low- and middle-income countries (WHO, 2018) due to their affordability. Sulfonamides act by competing with the precursor of dihydropteroate (DHP) synthase, p-aminobenzoic acid (PABA), which is crucial for folate biosynthesis. The folate pathway inhibition in turn blocks the bacterial purine synthesis for DNA production (Ovung & Bhattacharyya, 2019). The modified DHP synthases encoded by *sul* genes bypass the inhibition caused by sulfonamides, enabling resistance to these antibiotics (Nie et al., 2024).

Sul4 shares only 34 % amino acid identity with the DHP synthases and confers high-level resistance towards sulfonamides in *Escherichia coli* transconjugants (Razavi et al., 2017) even with one amino acid substitution (Peng et al., 2023). The near genetic context of *sul4* including integron integrase, a truncated *fol*K gene related to the folate biosynthesis pathway, and the ISCR element has already been confirmed by at least two studies (Peng et al., 2023; Razavi et al., 2017). However, the broader context and host range of *sul4* remain unclear because the initial discovery of the gene as well as the subsequent retrospective surveys relied on culture-independent methods (Berglund et al., 2023; Hutinel et al., 2022; Razavi et al., 2017). To our knowledge, the first, and currently only, description of *sul4* with confirmed sulfonamide resistance phenotype in a known bacterial host species was in a multidrug-resistant isolate of *Salmonella enterica* in 2023 (Peng et al., 2023). This discovery provides insight into the *sul4* gene’s potential to spread among clinically significant strains.

Despite their extensive use over the decades, new sequence variants of *sul* genes have emerged rarely, especially compared to beta-lactamase class ARGs. However, the latest observations of *sul4* suggest that there is a selection pressure for this gene to emerge and spread around the world (Berglund et al., 2023; Hutinel et al., 2022; Peng et al., 2023; Razavi et al., 2017). Still, the variety of the hosts and mobility mechanisms must be investigated to be able to understand the risk posed by *sul4* for the efficacy of sulfonamide antibiotics. In this study, we aimed to reveal the different host bacteria and genetic contexts of *sul4* found in influent and effluent wastewater as well as dried sludge. Wastewater collects fecal material and gut microbiota from entire populations linked to the sewer system. Furthermore, the communal wastewater resistome, which encompasses all detected ARGs, seems to reflect the resistance patterns observed in clinical reports and capture the spatio-temporal variations in resistance trends across populations (Karkman et al., 2020; W. Li et al., 2022; Pärnänen et al., 2019).

Different compartments of WWTP show differences in the diversity and composition of resistomes and this is linked to the changes in the microbiomes (Bengtsson-Palme et al., 2016; Ju et al., 2019). Activated sludge holds a central role in wastewater treatment as it enables the micro-organisms to work for nutrient removal by different biochemical processes (Nie et al., 2024). Contradictory results have been presented regarding the removal versus enrichment of ARGs and for instance integron integrase encoding genes during the treatment process (Bengtsson-Palme et al., 2016; Berglund et al., 2023; Guo et al., 2017). However, these genes are transcribed more prominently in the WWTP outlet than in the incoming wastewater, reflecting bacterial responses to environmental stressors such as antimicrobial compounds and rapidly changing conditions during the treatment process (Ju et al., 2019). While an effective wastewater treatment system removes most bacteria and the ARGs with them, some ARGs may become enriched during the process due to the proliferation of their host bacteria (Bengtsson-Palme et al., 2016; Guo et al., 2017) and even end up in the wastewater treatment plant (WWTP) discharge (Che et al., 2019).

Although metagenomic sequencing in wastewater detects a wide variety of ARGs, this method, particularly short-read sequencing, often fails to reveal how these genes are carried, transmitted, or their origin. Yet, this information corresponding to each ARG’s risk and significance in the overall resistance burden is crucial, especially in the case of novel emerging ARGs (Inda-Díaz et al., 2023; Nielsen et al., 2022). Connecting functional genes such as ARGs and mobile genetic elements to their carrier bacteria remains a challenge of the metagenomic approach (Abramova et al., 2024; Kerkvliet et al., 2024). Here we used PacBio long-read metagenomic sequencing with read lengths extending well over typical ARG lengths to enable reliable investigation of *sul4* gene contexts. Additionally, we took advantage of the unique bacterial methylation patterns to resolve the bacterial hosts for *sul4* in the complex wastewater microbial communities. DNA methylation catalyzed by MTases is the most important epigenetic modification mechanism in bacteria (Seong et al., 2021). Due to the bacterial differences in MTase presence, species-specific methylation profiles have been used to resolve genetic material of shared origin also before (Beaulaurier et al., 2018). Current algorithms for binning metagenome-assembled genomes (MAGs) using sequence composition and differential coverage may struggle to detect strain-level differences or, for example, link plasmid sequences with varying coverages to their associated host genomes (Beaulaurier et al., 2018). Applying methylation signals for linking contigs together according to their source organism can help to overcome these limitations.

## Materials and methods

### Sample collection

Composite samples from influent and effluent wastewater and dried sludge were collected on three days in 2019 (18.3.2019, 20.3.2019, 26.3.2019) at one of the two wastewater treatment plants responsible for processing wastewater in the Helsinki metropolitan area in Finland. The population of this area is about 1,6 million. In 2019, the sampled WWTP received 107 million m³ of influent wastewater, accounting for 71.8% of the total influent volume in the Helsinki Metropolitan area. This was due to the other WWTP being phased out and replaced by a new facility in the following years. The sampled treatment plant received wastewater from households and industry. In 2019 the latter made approximately 5 % of the influent when counting together both WWTPs at the Helsinki metropolitan area. Rainwater runoff is also directed to the WWTP. In the sampled WWTP, nearly 65,000 kg of sludge was produced by the treatment process in 2019. 93% of the dried sludge was delivered to several companies for further processing by composting-based methods, ultimately resulting in products suitable for agricultural or landscaping use through composting. Part of the sludge composting runoff was pumped back to the WWTP.

### DNA extraction, sequencing, and preliminary read processing

Wastewater samples were filtered through a 0.22 µm filter, and DNA was extracted from the filter. The Qiagen DNeasy PowerWater kit was used with bead beating for extracting the DNA, which was sent to the Institute of Biotechnology, University of Helsinki for library preparation and PacBio long-read metagenomic sequencing. PacBio Sequel II instrument was used for Single Molecule Real-Time (SMRT) sequencing with 12 SMRT cells. High Fidelity (HiFi) reads with kinetics information were prepared using PacBio BAM toolkit programs (https://github.com/PacificBiosciences/pbtk) to allow methylation detection while reads without the tags were used for metagenomic assembly. Additionally, paired-end short-reads were obtained using Illumina Novaseq. Short-read data was used for read recruitment for metagenome-assembled genomes (MAGs) and the bacterial diversity analysis by Metaphlan4 (Blanco-Míguez et al., 2023). These data were analyzed and visualized using phyloseq (McMurdie & Holmes, 2013), vegan (Oksanen et al., 2020), and ggplot2 (Wickham, 2016) in R (v4.4.0) (R Core Team, 2023) and RStudio (v2024.12.0+467.pro1) (RStudio Team. Inc. Boston, 2016) (Supplementary Methods).

### Metagenomic assembly and identification of *sul4* reads and contigs

To improve the reliability of the assemblies, two assemblers were used for the metagenomic assembly of the wastewater samples, hifiasm-meta (v0.18.0) (Feng et al., 2022) and metaFlye (v2.9.3) (Kolmogorov et al., 2020). The first mentioned was run with the default options and the latter with the additional parameter --min-overlap 4000. Reads encoding *sul4* were extracted using a custom Snakemake (Mölder et al., 2021) workflow, which included steps for mapping the reads to an indexed *sul4* reference using pbmm2 align (v1.9.0) (https://github.com/PacificBiosciences/pbmm2) and converting the mapped reads from bam to fasta files with bam2fasta (v1.0.0) (https://github.com/PacificBiosciences/pbtk). The *sul4* contigs were extracted from the sample assemblies using BLAST (Camacho et al., 2009).

### Collection of *sul4* host sequences from public databases

The *sul4* reference gene (NG_056174.1) was used to explore the matching sequences in the National Center for Biotechnology Information (NCBI) and Integrated Microbial Genomes and Microbiomes (IMG) databases. Sequences with an alignment length greater than 500 bp, relative to the reference gene (864 bp), were included in the downstream analysis, yielding 11 hits. Data collection was completed by the 30^th^ of April 2024.

### Methylation analysis for *sul4* contigs and wider set of contigs

Methylation analysis was run for all 12 *sul4* contigs sequenced and assembled here. More detailed methods for methylation analysis are found in the Supplementary Methods. Briefly, PacBio reads with kinetics tags were mapped against the assembled contigs, and the kinetics information in the mapping reads were summarized by ipdSummary (v3.0) (https://github.com/PacificBiosciences/kineticsTools/). MultiMotifMaker algorithm (Li et al., 2020) was run to predict the sequence motifs surrounding the methylated bases. The methylation motifs of types m6A and m4C were explored and visualized using pheatmap (v 1.0.12) (Kolde, 2019) in R (R Core Team, 2023) and RStudio (v2024.12.0+467.pro1) (RStudio Team. Inc. Boston, 2016).

To explore contigs with similar methylation profiles, methylation analysis was run for a wider set of contigs from all wastewater samples. However, due to the resource-intensive nature of *de novo* motif prediction, only a subset of the contigs from the entire assembly could be analyzed for methylation profiles (Supplementary Figure 1). During the exploration of methylation profiles similar to *sul4*-containing contigs, those matching only the G<u>A</u>TC (m6A) motif were excluded. This approach was chosen because of the high prevalence of this motif owing to its regulatory roles among various bacterial taxa (Seong et al., 2021) could overshadow the patterns of other motifs (Beaulaurier et al., 2019; Roberts et al., 2023; Sánchez-Romero & Casadesús, 2020; Seong et al., 2021).

### Gene annotations of *sul4* sequences

The bacterial genome annotation tool Bakta (v1.7.0) (Schwengers et al., 2021) was run for all *sul4* sequences including those sequenced here and those obtained from public sequence databases to get the coordinates for *sul4* genes in these sequences. Sequence region 10 kbp up and downstream *sul4* gene were extracted by custom script applying SeqKit tools (v2.5.1) (Shen et al., 2016). After dereplication of *sul4* sequences sharing above 90 % similarity using CD-HIT (v 4.8.1) (Fu et al., 2012), Proovframe (v0.9.8) (Hackl et al., 2021) applying Diamond (v2.0.15) (Buchfink et al., 2021) was used to identify and correct indels causing frameshifts in coding sequences and affecting gene prediction. Finally, the remaining 34 ∼20 kbp *sul4* flanking sequences were visualized using clinker (v 0.0.27) (Gilchrist & Chooi, 2021).

### Investigation of *sul4* module genes

Integron integrase genes from *sul4* sequences were extracted and aligned, along with five reference integrase genes from NCBI, using MAFFT (v7.505) (Katoh et al., 2002). Half of the *sul4* sequences (n=17) were included in the integrase gene sequence similarity analysis, as in some cases, the contig or read was cut at the gene site. The reference genes were selected based on their similarity to integrase genes from different branches of the initial tree, identified through a BLAST search against the NCBI database. Additionally, a reference gene of class 1 integron integrase encoded by *Escherichia coli* EC261 (AP027420.1) was included in the analysis. The multiple sequence alignment was rooted with site-specific recombinase genes *xerC* (NC_000913.3:3996287-3997183, GeneID: 948355) and *xerD* (NC_000913.3:3038847-3039743, GeneID: 947362) in *Escherichia coli* str. K-12 substr. MG1655. Like integron integrases, these genes are tyrosine recombinases that mediate site-specific DNA recombination (M. C. Liu et al., 2022). The phylogenetic tree was drawn using RAxML (v 8.2.12) (Stamatakis, 2014) with model GTRGAMMA and visualized with iTOL (Letunic & Bork, 2021). The *sul4* module ISCR sequences were extracted through the detection of *ori*IS and *ter*IS sequences specific to these elements (Yuan et al., 2024) (Supplementary Methods). HattCi (v1.0b) (Pereira et al., 2016) was used to investigate the *attC* sites as putative integron-associated target sequences for the ISCR elements. The relatedness of the ISCR sequences present in our *sul4* modules compared to the known ISCR genes and canonical IS91 family members (Yuan et al., 2024) was studied using multiple sequence alignment with Clustal Omega (Madeira et al., 2024). The analysis of the phage-related genes involving geNomad (Camargo et al., 2023) and iPHoP (Roux et al., 2023) is found in the Supplementary Methods.

### Plasmid prediction

The plasmid contigs from wastewater metagenomes were predicted using two programs plasX (Yu et al., 2024) and geNomad (v1.8.0) (Camargo et al., 2023) to determine whether the detected *sul4* genes were located on plasmids and to filter data for the broader methylation analysis, for which only putative plasmid contigs longer than 30 kb were considered (See section ‘Methylation analysis’). For plasX, Anvi’o (v8) (Murat Eren et al., 2021) was used to generate contigs database with functional annotations from COG (database version COG_2014) and Pfam (database version Pfam_v32) databases before running the plasX predictions. The geNomad program was run with the default parameters.

### Curation of metagenome-assembled genomes (MAG) for *sul4* carriers

MAGs containing gene *sul4* were curated through manual binning using Anvi’o (v8) (Murat Eren et al., 2021) integrated into a custom Snakemake (Mölder et al., 2021) workflow. The whole process was done separately for the two different assemblies generated by hifiasm-meta and metaFlye. A more detailed description of the programs and analysis scripts used is found in the Supplementary Methods. In brief, functional gene annotations including ARG annotations with ResFinder (v4.2.3) (Bortolaia et al., 2020) were done for contigs databases built according to Anvi’o workflow. Illumina short-reads were recruited to gather the reads aligning to the contigs. MAGs for *sul4* carrying contigs were manually refined within the Anvi’o interactive interface visualizing each metagenomic sample at a time. Quality check with CheckM2 (Chklovski et al., 2023), as well as taxonomical identification with GTDB-Tk (database version 220) (Parks et al., 2022), was done for the manually refined *sul4* MAGs. MAGs fulfilling the criteria of completion and redundancy values of 50 and 10 respectively were kept for the preliminary analysis. Unique high and middle-quality MAGs were described in more detail.

### Analysis of non-*sul4* contigs and MAGs

The non-*sul4* contigs with similar methylation profiles to some of the *sul4* contigs were analyzed similarly to the *sul4* contigs regarding plasmid prediction and the analysis of the *sul4* flanking region. One of these five contigs, which showed a similar phage sequence region to some of the gene *sul4* encoding contigs, was investigated in more detail and visualized alongside the *sul4* contigs. As these five contigs originated from two different samples (SLU1 and SLU2) they were combined sample-wise into two separate MAGs of two and three contigs. The resulting MAGs were assessed for quality using CheckM2 (Chklovski et al., 2023), and their taxonomic identity was determined with GTDB-Tk (Parks et al., 2022).

### Data availability

The sequence data will be available under accession number PRJEB83306 in the European Nucleotide Archive (ENA). The MAG sequences are available in Figshare under the doi X (TBA). The analysis scripts are shared in the publicly available repository at https://github.com/melinamarkkanen/sul4_project.

## Results

### *sul4* was detected in all but one of the wastewater samples including influent, effluent, and dried sludge

Most of the metagenomic reads mapping to *sul4* originated from sludge samples while reads from influent and effluent were fewer (Table 1). Moreover, *sul4*-containing contigs could be obtained only from sludge metagenomic assemblies (Table 2), where *sul4* was more commonly detected and the bacterial diversity was decreased in contrast to the influent (Supplementary Figure 2). The average length of *sul4* contigs ranged from 8 636 to 1 884 747 bp (Table 2). Four and seven contigs with *sul4* were constructed using the two different assemblers hifiasm and metaFlye, respectively (Table 2). According to the plasmid prediction analysis with plasX and geNomad, only one of the *sul4* contigs (contig_s43139.ctg070500l) showed plasmid-like properties and this was identified solely by, geNomad (Table 2). However, this contig was notably shorter than the other *sul4* contigs (8 636 bp), which might have affected the plasmid prediction algorithms’ abilities to correctly identify plasmid sequences.

**Table 1.** Reads mapping to reference gene *sul4* (NG_056174.1)

| Sample code | Sample | Reads mapping to *sul4* |
| --- | --- | --- |
| INF1 | influent | 1 |
| INF2 | influent | 0 |
| INF3 | influent | 3 |
| EFF1 | effluent | 3 |
| EFF2 | effluent | 1 |
| EFF3 | effluent | 1 |
| SLU1 | sludge | 23 |
| SLU2 | sludge | 34 |
| SLU3 | sludge | 8 |

**Table 2.** Assembly statistics of *sul4* contigs. The plasX scores below 1×10^-3^ are denoted as 0.

| Sample | Assembler | No. of contigs | Contig names | Contig length (bp) | Score for plasmid plasX / geNomad |
| --- | --- | --- | --- | --- | --- |
| SLU1 | hifiasm | 1 | contig_s1.ctg000044l | 1 884 747 | 0 / 0 |
| SLU2 | hifiasm | 1 | contig_s3.ctg001711l | 109 015 | 0 / 0 |
| SLU3 | hifiasm | 2 | contig_s3566.ctg005183l<br>contig_s43139.ctg070500l | 314 103<br>8 636 | 0 / 0<br>0.003 / 0.97 |
| SLU1 | metaFlye | 2 | contig_13571<br>contig_73809 | 525 812<br>143 767 | 0 / 0<br>0 / 0 |
| SLU2 | metaFlye | 4 | contig_14681<br>contig_49547<br>contig_49550<br>contig_49551<br>contig_64719 | 50 817<br>19 369<br>38 474<br>396 357<br>793 955 | 0 / 0<br>0.007 / 0<br>0 / 0<br>0 / 0<br>0 / 0 |
| SLU3 | metaFlye | 1 | contig_65443 | 131 210 | 0 / 0 |

### The conserved module containing *sul4*, truncated *fol*K, and an ISCR element was found in diverse genetic contexts

A highly conserved gene region flanking the *sul4* (hereafter *sul4* module) was identified in all *sul4* reads and contigs from our wastewater samples. Furthermore, with a few exceptions where the sequence was cut probably leaving out some of the genes, an integron integrase was always linked to the *sul4* module (Figure 1). The core structure of the *sul4* module consisted of a partial *fol*K gene, *sul4,* and an ISCR element found downstream of *sul4* (Figure 1). To further investigate the conservation, variation, and potential role of this module in mobilizing *sul4*, we expanded our analysis to include sequences encoding *sul4* found in public sequence databases. Screening sequence data in the National Center for Biotechnology Information (NCBI) and Integrated Microbial Genomes and Metagenomes (IMG) with reference gene *sul4* (NG_056174.1) resulted in 11 different hits with good match (length above 500 bp of the *sul4* gene length 864 bp). These matches included the already published *sul4* sequences by Razavi and colleagues (NG_056174.1, MG649394.1, MG649402.1, uncultivated bacteria) (Razavi et al., 2017) and Peng and colleagues (CP120671.1, *Salmonella enterica*) (Peng et al., 2023) as well as those sequences without previous description regarding their *sul4* carriage (Table 3). A highly conserved *sul4* module was present in all of these sequences. We also noted the *sul4* resembling genes published by Shindoh and colleagues (Shindoh et al., 2023). However, we excluded them from our analysis as they showed only minor similarity to the *sul4* reference gene (NG_056174.1).

**Figure 1.**
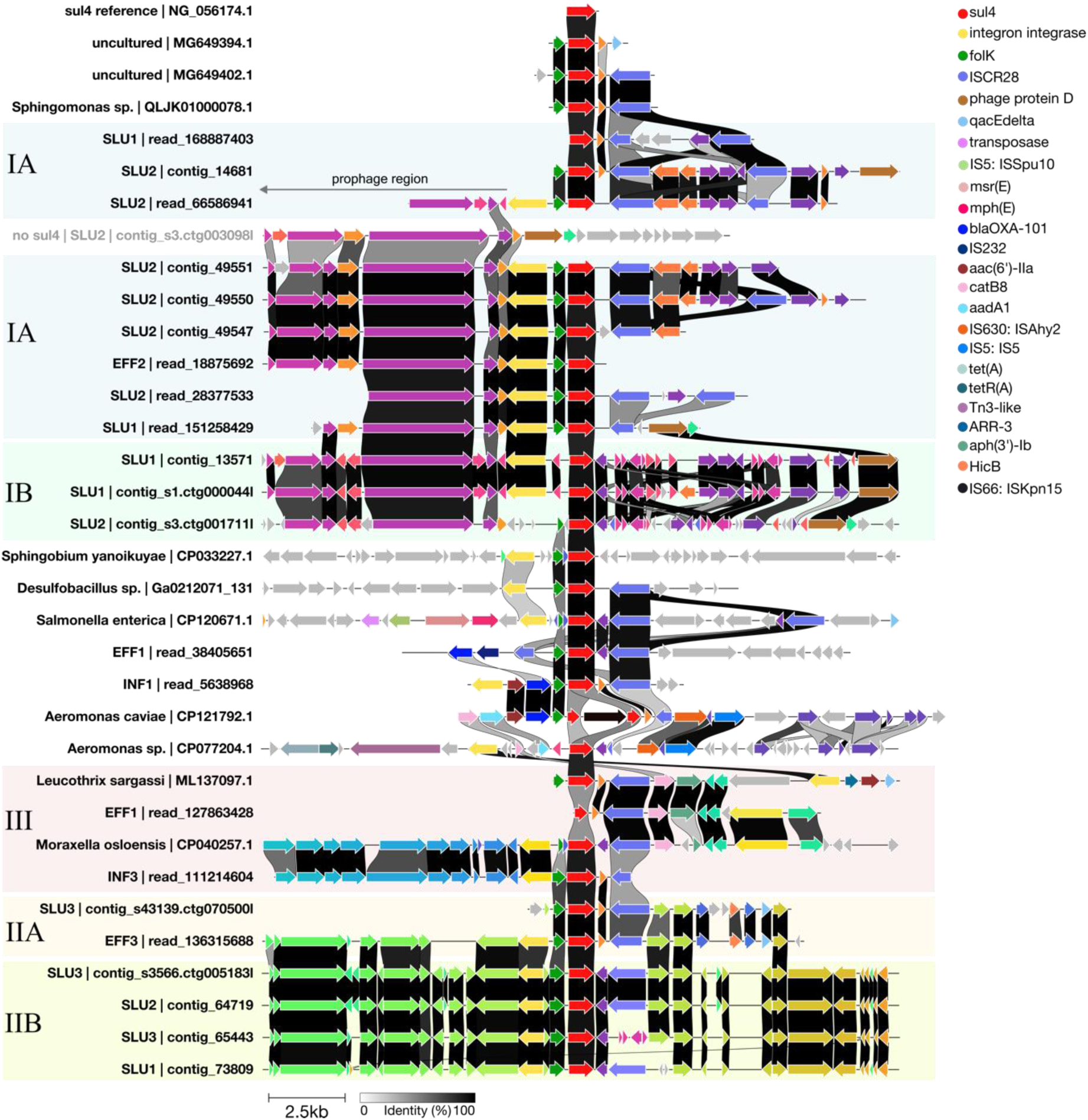
Genetic contexts of 10 kb upstream and downstream of *sul4* gene in long-reads and contigs sequenced here and *sul4* accessions published previously. Visualization is done using Clinker based on Bakta annotations. The alignments are centralized to the *sul4* gene colored red. ARGs and key mobile genetic elements are color-coded and explained in the legend on the up-right. Other gene annotations without a specified color label are displayed using the default colors by Clinker, which assigns consistent colors to genes with similar sequences. Vertical links indicate the similarity of genes between different sequences (black = identical; white = no similarity).

**Table 3.**
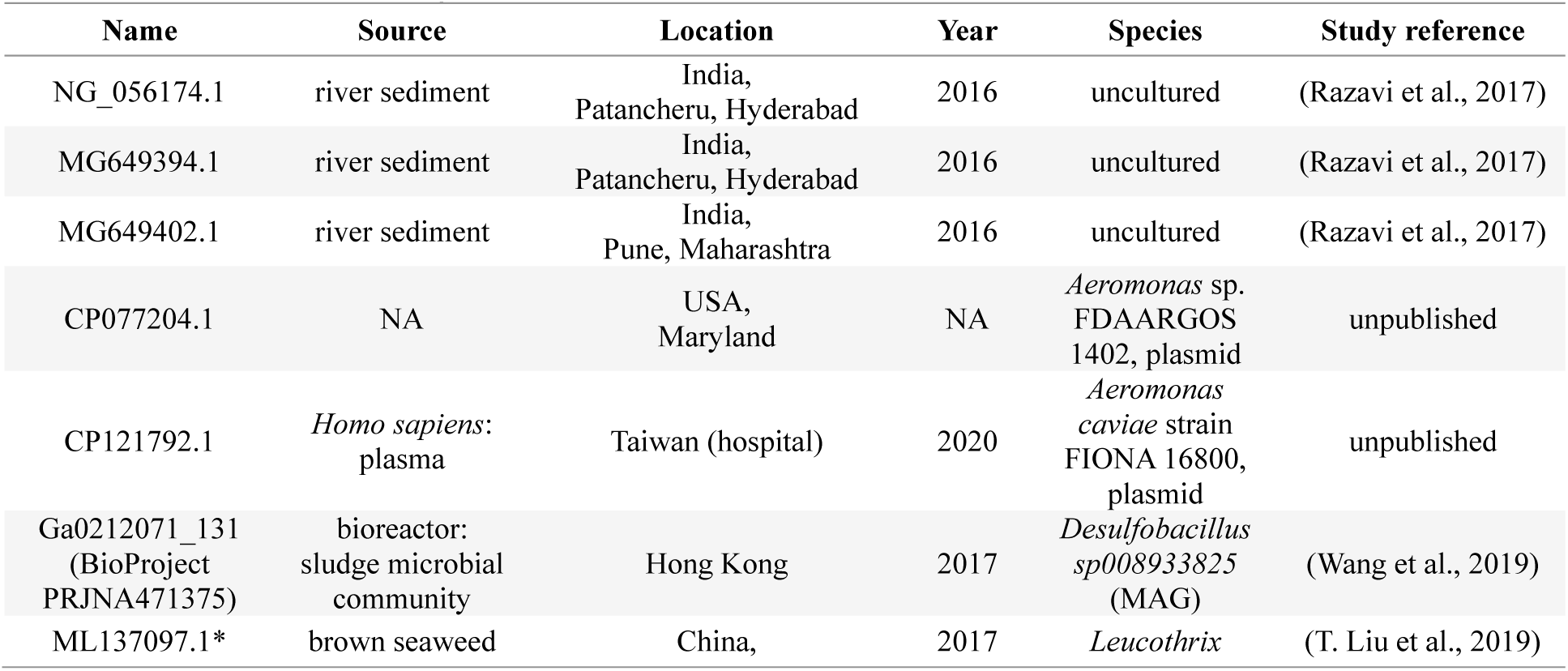
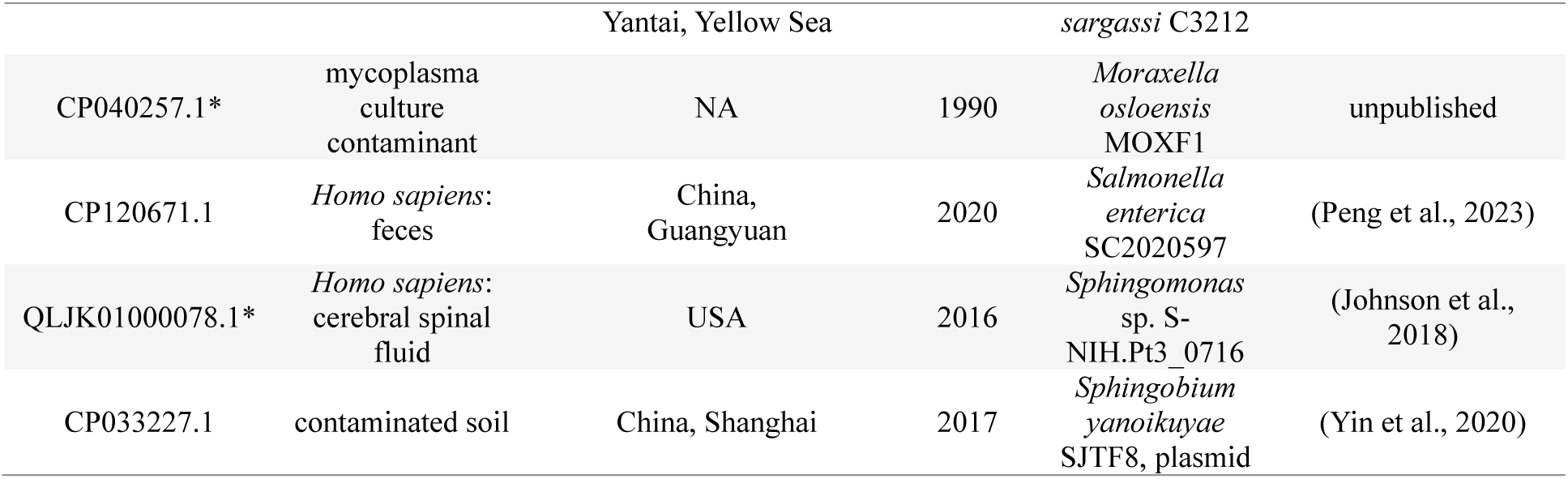
Reference sequences for data visualized in Figure 1. Sequences marked with an asterisk denote suppressed accessions in the RefSeq database due to putative contamination or unconfirmed source organism.

The source environment of *sul4* sequences from databases varied greatly ranging from anthropogenically contaminated river sediment to seashore marine seaweed, as well as clinical isolates of human origin and microbes from wastewater bioreactors (Table 3). These species included opportunistic human pathogens *Salmonella enterica*, *Aeromonas* spp., as well as *Sphingomonas* sp. and *Moraxella osloensis* (Table 3), which also contain pathogenic strains, though less frequently. The remaining *sul4* carrier bacteria from sequence databases included the species *Leucothrix sargassi*, *Sphingobium yanoikuyae*, and *Desulfobacillus sp008933825.* However, as an important note, three of the NCBI reference sequence (RefSeq) accessions for *sul4* carrier bacteria; GCF_005518075.1 for *Moraxella osloensis* MOXF1 (CP040257.1), GCF_004010795.1 for *Leucothrix sargassi* C3212 (ML137097.1), and GCF_003950715.1 for *Sphingomonas* sp. S-NIH.Pt3_0716 (QLJK01000078.1) had been suppressed as they did not meet the criteria of RefSeq data. For instance, for the *M. osloensis* strain MOXF1, the assembly coverage was 70.84% which indicates that ∼30% of this genome is not *M. osloensis*. Instead, for *L. sargassi* C3212, ∼30% of the genome is a good match for *Moraxella tetraodonis* referring to contamination in this genome. Finally, the RefSeq accession for *Sphingomonas* sp. S-NIH.Pt3_0716 had been suppressed as the source organism could not be confirmed. This information was considered when evaluating the reliability of these three genomes. Geographically, the *sul4* sequences found in sequence databases originated from different parts of the world with an emphasis on Asian countries (Table 3).

Wider genetic contexts surrounding *sul4*, including sequences 10 kb upstream and downstream of the *sul4* gene, were analyzed and visualized for all *sul4* sequences (Figure 1, Table 3). The genetic contexts beyond the more conserved *sul4* modules exhibited greater variation and could be classified into three major groups (I-III) by visual inspection, each showing high similarity in gene content (Figure 1). More specifically, groups I and II contained sequences with moderate similarity, which led to their division into subgroups IA-B and IIA-B. The remaining wider genetic contexts were either unique or too short to be described as members of any of the groups I-III (Figure 1).

### Subtle sequence variations in *sul4* modules and adjacent integron integrases were observed across the different contexts

Although the core structure of *sul4* was highly conserved among all investigated sequences, slight differences were also noted (Figure 1). First, the comparison of the integrase sequences upstream *sul4* module resulted in roughly 6 clusters (Figure 2). These clusters were accompanied by a reference integrase gene from previously published sequences (Figure 2).

**Figure 2.**
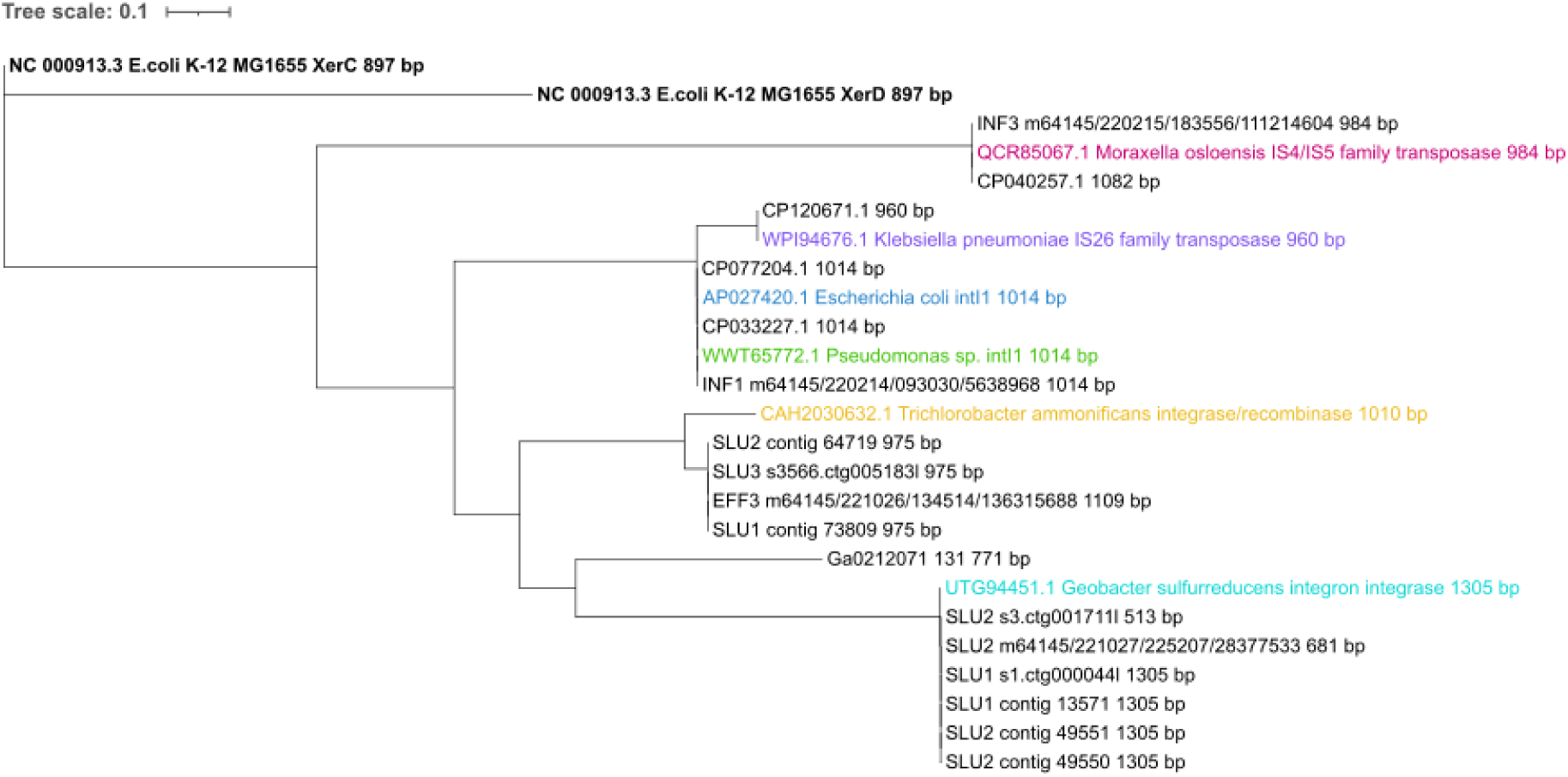
Phylogenetic tree of integron integrase sequences in *sul4* sequences. The tree was rooted by XerC and XerD recombinase encoding genes in type strain *E. coli* K-12 MG1655 (bold). Multiple sequence alignment was run using MAFFT and the tree was computed with RAxML. Interactive Tree of Life (iTOL) was used for visualization. The other reference integrase genes are highlighted with colors other than black.

An exception to this was the *Desulfobacillus sp008933825* (Ga0212071_131) encoding integrase gene which was shorter and most different from any other of the integrases (Figure 2). Integrase genes from *sul4* sequences, classified as group I based on their broader genetic context (Figure 1), were grouped into a single cluster (Figure 2). This cluster was represented by the reference integrase gene from *Geobacter sulfurreducens* (Figure 2). Integrases of *sul4* sequences of group II (Figure 1) showed the closest but not perfect similarity with the *Trichlorobacter ammonificans* integrase reference gene (Figure 2). A third cluster was formed by integrases from one influent wastewater read and the *M. osloensis* (CP040257.1) strain, both belonging to group III (Figure 1). This cluster was represented by a reference integrase gene by another *M. osloensis* strain (Figure 2).

The separate cluster of the integrases in *Aeromonas* spp. (CP077204.1) and *Sphingobium yanoikuyae* (CP033227.1) were accompanied by one influent read and reference integrase genes of class 1 integron integrase (*intI1*) found in *Pseudomonas* sp. and *E. coli* (Figure 2). According to a BLAST search against the NCBI database, identical *intI1* genes are also found in many other gram-negative opportunistic human pathogens of class Gammaproteobacteria (data not shown). The integrase gene in *sul4* host *S. enterica* (CP120671.1) showed high similarity with the reference integrase gene hosted by gram-negative opportunistic pathogens such as *Klebsiella pneumoniae*. These two integrases showed the highest similarity with the *intI1* cluster.

Additional ARG properties beyond *sul4* were detected in part of the sequences. Specific to *Sphingobium yanoikuyae* strain (CP033227.1), a putative aminoglycoside riboswitch, linked to the integron *attI* site, was identified between the integrase and *sul4* (Figure 1), while some of the ARGs *aac(6’)-IIa, bla*OXA-101*, catB8* and *aadA1* were found to be part of the *sul4*-associated integron cassettes in one influent wastewater read and *Aeromonas* spp. plasmid sequences (Figure 1). Specific to the group III sequences, also secondary integrases with ARG arrays that were located more distant downstream to *sul4* were observed (Figure 1). Altogether, the variation observed in the arrangement and phylogeny of these integrase sequences, extending beyond those of class 1 integrons, suggests their enduring role in linking and potentially mobilizing *sul4* across different contexts.

According to our phylogenetic analysis, the ISCR elements present in nearly all *sul4* modules investigated here, across various wider contexts, were identified as ISCR28 (Yuan et al., 2024) as they were dissimilar to the other ISCR elements, including ISCR-20, and canonical IS91-like genes (Supplementary Figure 3). We detected both *ori*IS and *ter*IS or at least *ori*IS sequences in all these ISCR28 elements (data not shown). Also, integron-associated *attC* sites were identified at the end or right after each ISCR28 gene (data not shown). As an important note, although in Figure 1 the gene annotations for ISCR28 in group IB as well as some of those in group IIB appear substantially distinct from the ISCR28 in the other sequences, our phylogenetic analysis confirmed that they are identical, differing only slightly in length (Supplementary Figure 3). This incorrect visualization was caused by the frameshift changes affecting gene annotation. Secondary ISCR genes further on the right-hand side were common to many group I sequences and *S. enterica* (CP120671.1) (Figure 1). Instead in one effluent read, an additional remnant of ISCR28 element was located also upstream of *sul4* next to the *bla*OXA-101 gene (Figure 1).

However, in three *sul4* modules, the IS element was dissimilar to ISCR28 and ISCR20: These elements, found in the two plasmids of *Aeromonas* (CP121792.1 and CP077204.1), and *Sphingobium yanoiukuyae* (CP033227.1) grouped into their separate clusters (Supplementary Figure 3). The BLAST search of the IS element of *S. yanoikuyae* (CP033227.1) revealed that part of this gene is found primarily in other members of *Sphingobium* and relative genera (data not shown). The IS element genes in the *Aeromonas* spp. *sul4* plasmids were in fact annotated as three genes: IS630-like ISAhy2 transposase, hypothetical protein-encoding gene, and IS5-like IS5 element. According to a BLAST search against the NCBI database similar gene regions are found in various other Gammaproteobacteria including opportunistic pathogenic genera such as *Pseudomonas, Salmonella, Klebsiella,* and *Enterobacter* (data not shown).

Taken together, ISCR28 was found in most of the *sul4* modules investigated. However, two other kinds of clearly distinct IS sequences at this location were found: One in the *sul4* carrying *Aeromonas* plasmids (and found in other typical opportunistic pathogen Gammaproteobacteria) and the other one hosted by a plasmid of *Sphingobium yanoikuyae* and found in closely related taxa.

Compared to the integrases and IS elements discussed above; fewer dissimilarities were detected for the gene *sul4* itself (Figure 1). One such being however, *Aeromonas caviae* FIONA 16800 (CP121792.1) where the *sul4* gene was cut into two parts by incorporated IS66-like IS*Kpn15* transposase gene (Figure 1).

### *sul4* modules were identified in various wider contexts, exhibiting minimal to moderate similarity to one another

Analysis with geNomad revealed that the *sul4* module in group I (both subgroups IA and IB) sequences including reads and contigs from effluent and sludge samples was located adjacent to a prophage sequence region (Supplementary Tables and Supplementary Results). Part of this region is visible in Figure 1 downstream integrase gene of group I sequences (genes on the left-hand-side of group I in Figure 1). These prophage sequences were identified to belong to class Caudoviricetes (Supplementary Tables and Supplementary Results).

The sequence regions examined in groups IIA and IIB were highly similar, with the main difference being the presence of type 2 toxin-antitoxin system HicA and HicB family toxin encoding genes downstream of ISCR28 in group IIA and their absence in group IIB (Figure 1). Group III was formed of both sequences of this study and those from the public sequence databases (Figure 1). The two reads from influent and effluent wastewater resembled the wider genetic context of *sul4* seen in *Moraxella osloensis* (CP040257.1) and *Leucothrix sargassi* C3212 (ML137097.1) (Figure 1). More specifically, the influent read aligned with the upstream and the read from effluent with the downstream sequence of *sul4*. The region further downstream *sul4* contained another integron cassette with *ARR-3*, *aac(6’)-Ib*, *qacEdelta1,* and *sul2* (data not shown). However, in *L. sargassi*, a gene encoding a protein of unknown function seemed to truncate the secondary integron cassette (Figure 1).

In addition to the groups described above, a miscellaneous set of genetic contexts sharing varying levels of similarity among each other was observed (Figure 1). These sequences included the recently described known *sul4* carrier *Salmonella enterica* (CP120671.1), and those found in public databases without previous reports on their *sul4* carriage; *Sphingobium yanoikuyae* (CP033227.1), *Sphingomonas* sp. (QLJK01000078.1), *Aeromonas* spp. plasmids of two different strains (CP077204.1 and CP121792.1) and *Desulfobacillus sp008933825* (Ga0212071_131). The two distinct *Aeromonas* strain plasmids carrying *sul4* showed similar genetic composition with each other and partially with one influent wastewater read. The remaining *sul4* sequences identified as *S. enterica*, *D. sp008933825, S. yanoikuyae*, and *Sphingomonas* sp. were all unique regarding the wider genetic context of *sul4* (Figure 1).

### Unique methylation profiles indicated the presence of distinct *sul4* hosts in wastewater

After finding the conserved *sul4* module in different wider contexts in our wastewater samples, we were interested in identifying the bacterial species carrying these sequences. For that, we exploited bacteria-specific methylation patterns of the *sul4* encoding contigs. On average six methylated motifs (ranging from 1 to 13) of types m6A and m4C were detected among the *sul4* contigs (Figure 3). One or more methylated motifs were detected for all but one *sul4* contig (contig_49547) for which no methylated motifs were identified (Figure 3). The most common methylated motif, detected in six *sul4* contigs, was G<u>A</u>TC. A distinctive cluster was formed of *sul4* contigs possessing the methylated motif G<u>A</u>TC together with at least one of the five other motifs, AGG<u>C</u>YG, GH<u>A</u>CNNNNTCC, GG<u>A</u>NNNNGTDC, CTCG<u>A</u>G, and RCCC<u>C</u>C (Figure 3). The two other *sul4* contigs with methylated motif G<u>A</u>TC (Figure 3) lacked any other motifs of this cluster. The remaining five *sul4* contigs had no methylated motifs in common with any of the other contigs (Figure 3). The methylation-based grouping of contigs into three categories: those with a uniform profile based on several motifs, those with G<u>A</u>TC but otherwise unique profile, and those with no consistent pattern (Figure 3) suggested that at least two distinct bacterial species were carrying the *sul4* gene in our wastewater samples.

**Figure 3:**
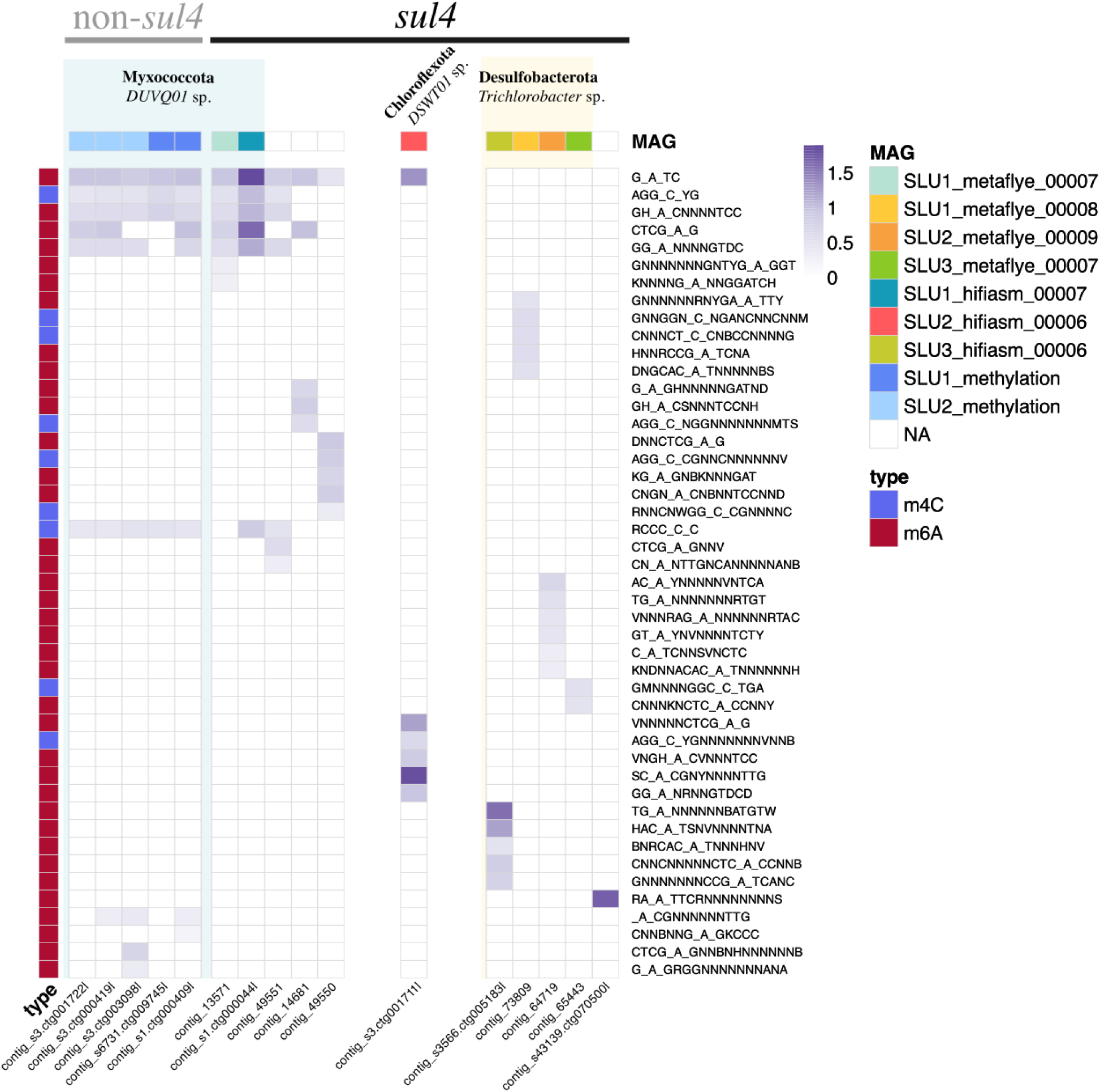
Methylation profiles of *sul4* contigs and contigs without *sul4*. Contigs are represented as columns and the methylation motifs are shown as rows on the right-hand side of the heatmap. The intensity of the purple color in the heatmap shows the fraction of the methylated sites for each of the motifs predicted by the MultiMotifMaker algorithm (Li et al., 2020). The modification type (m6A or m4C) is indicated by color coding on the left-hand side. The top row annotation indicates the MAG in which each contig is included, if applicable, and the taxa assigned to the MAGs are shown in the text.

To test the discriminatory power of methylation signatures for binning species-specific contigs and to find contigs of common origin species with *sul4* contigs, a wider set of contigs was examined in search of similar methylated motifs displayed by the 12 *sul4* contigs. For that, methylation profiles of large chromosomal contigs of above 200 kbp and putative plasmid contigs of above 30 kbp in the length of the hifiasm assembly were explored. Such stringent filtering was necessary due to the computationally intensive *de novo* methylation motif finder algorithm, which could only be run on a subset of the entire metagenomic dataset. Applying these criteria resulted in a small subset of contigs from the full assemblies (1.38%) being retained for the methylation analysis (Supplementary Figure 1). The proportion of large contigs studied for methylation was highlighted in sludge samples while the number of putative plasmid contigs was higher in influent and effluent samples (Supplementary Figure 1). Using this approach, we identified five non-*sul4* contigs with methylation profiles matching those of the *sul4* cluster, all exhibiting virtually uniform profiles (Figure 3). We therefore hypothesized that these contigs originated from genomes of the same bacterial species. To confirm this and further investigate the taxonomical identity of these sequences, we next focused on generating metagenome-assembled genomes (MAGs) for these contigs.

### The presence of three distinct bacterial hosts for *sul4* in wastewater sludge was proven by metagenome-assembled genomes

Metagenome-assembled genomes (MAGs) were generated to verify the distinct *sul4* host taxa in wastewater, that were indicated by the methylation profiles. Binning of *sul4* MAGs from three sludge samples and for assemblies from both assemblers was done using Anvi’o (Eren et al., 2020) (Table 4). Altogether seven MAGs with *sul4* were generated. Two out of them (SLU2_metaflye_00009 and SLU1_metaflye_00007) met the criteria of a high-quality MAG (> 90% completeness, < 5% redundancy) defined by Genome Standards Consortium (GSC) and verified by CheckM2 (Table 4). The remaining five MAGs were medium-quality with 50–90% completeness and less than 10% contamination (Table 4). Both high-quality *sul4* MAGs originated from metaFlye assemblies of two sludge samples (SLU1 and SLU2) and represented genomes from two distinct bacterial phyla; Desulfobacterota and Myxococcota (Table 4). More specifically, according to the taxonomic assignment by GTDB-Tk, the Desulfobacterota MAG matched with the reference genome GCF_933509905.1 of *Trichlorobacter sp933509905* (nomenclature in NCBI: *Trichlorobacter ammonificans* G1) belonging to the class Desulfuromonadia (Table 4). Similarly, the Myxococcota MAG matched with reference genome GCA_012840965.1 assigned to species *DUVQ01 sp012840965* of class UBA9042 (Table 4).

**Table 4.**
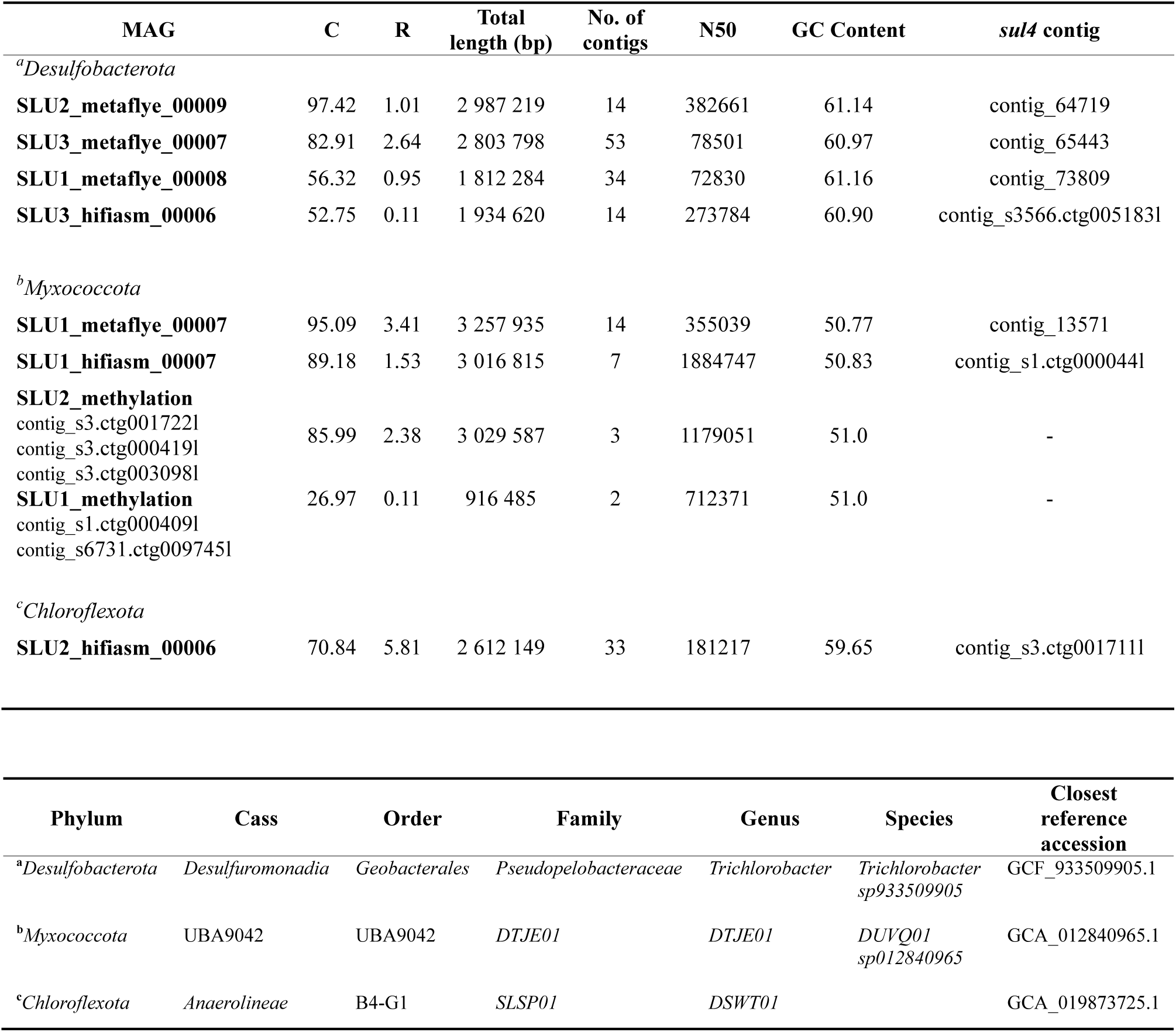
Manual binning of *sul4* MAGs and Myxococcota MAGs without *sul4* generated based on contigs with shared methylation profiles. Letters C and R refer to completeness and redundancy values estimated by CheckM2.

All except one (SLU2_hifiasm_00006) of the remaining medium-quality *sul4*-MAGs matched these two reference genomes and were found from two sludge samples (SLU1 and SLU3) (Table 4). Lastly, the medium-quality MAG SLU2_hifiasm_00006, distinct from all other MAGs, exhibited the closest match to a genome from the *DSWT01* genus of the Chloroflexota phylum (Table 4).

### Methylation-based binning revealed Myxococcota *DUVQ01 sp012840965* genomes lacking *sul4* gene

To confirm the identity of the non-*sul4*-containing contigs exhibiting similar methylation patterns as the *sul4*-containing cluster contigs in Figure 3, these contigs were binned together according to their origin sample (SLU1 or SLU2) resulting in MAGs SLU2_methylation and SLU1_methylation (Table 4). Similarly to *sul4* MAGs SLU1_metaflye_00007 and SLU1_hifiasm_00007, these MAGs were assigned as Myxococcota species *DUVQ01 sp012840965* (Table 4). While SLU2_methylation MAG was of medium-quality, showing 86.0 % completeness and 2.4 % redundancy values, SLU1_methylation was less complete (27.0 % completeness and 0.11 % redundancy) due to its short length. Hence, this bin was left out of further analyses (Table 4). Altogether the taxonomical identity of all Myxococcota MAGs generated for both *sul4* and non-*sul4* were in line with the initial methylation-based grouping of these contigs (Figure 3). This allowed us to investigate the differences seen in the genomes of these species with and without *sul4* more in detail (Figure 1). Interestingly, phage sequence regions similar to those contigs belonging to Myxococcota MAGs with *sul4* were observed also in those not carrying *sul4* (Figure 1).

## Discussion

In this study, we described several novel genetic contexts and host species for the recently discovered sulfonamide resistance gene variant *sul4* by the application of long-read metagenomic sequencing and unique methylation signatures. The three *sul4* hosts detected in wastewater samples were *DUVQ01 sp012840965* of Myxococcota phylum*, Trichlorobacter sp933509905* (nomenclature in NCBI: *Trichlorobacter ammonificans* G1) of Desulfobacterota phylum and species of genus *DSWT01* of Chloroflexota phylum. Moreover, by exploring data available in public databases, we identified additional *sul4*-carrying bacteria from various ecological niches and taxonomic lineages such as genera *Aeromonas, Moraxella, Sphingobium,* and *Desulfobacillus*, on top of the previously described *sul4*-carrying *Salmonella enterica* (Peng et al., 2023).

The core structure of the sequence region flanking *sul4* gene (*sul4* module) was highly conserved among different *sul4* hosts examined here and those described by previous research (Peng et al., 2023; Razavi et al., 2017) suggesting this module’s importance in the dissemination of *sul4* across bacterial lineages. The detection of the integron integrase gene in all studied *sul4* contexts (those where the sequence was intact at this site) increased the evidence for its speculated roles in the *sul4* gene expression regulation and mobilization (Peng et al., 2023; Razavi et al., 2017). Previous literature on integron-mediated spread of ARGs, including *sul4*, has focused on class 1 integrons (Gillings, 2017; Razavi et al., 2017) due to their substantial role in accelerating the antibiotic resistance crisis (Loot et al., 2024), as they can carry resistance gene cassettes among clinically important bacteria and between these and environmentally sourced species. As a putative example of the latter case, we found *sul4* in association with *intI1* in an environmental species *Sphingobium yanoikuyae* in a similar manner to *Pseudomonas* sp., *Aeromonas* sp., and other gram-negative opportunistic pathogens. This could support the hypothesis of class 1 integron-mediated movement of ARGs between environmental and clinically significant bacteria. As *intI1* linked *sul4* modules were present also in our wastewater data, within a genetic context similar to those in *Aeromonas* spp. this seems to represent a somewhat common gene arrangement.

However, according to our results also other less-studied types of integrons, typically originating from environmental settings, can act as key contributors in *sul4* dissemination; Our analysis revealed four other than class 1 integron-associated types of integrase genes linked to the *sul4* module. These integrase lineages mostly aligned with their broader context sequences, and host bacteria, suggesting species-specific adaptations. One of these was the integrases of our *DUVQ01 sp012840965* Myxococcota MAGs. Noteworthy, the closest reference gene matching this integrase was hosted by *Geobacter* species (phylum Desulfobacterota) and not species of Myxococcota phylum as anticipated. This was likely due to the unexplored diversity of the newly established phylum Myxococcota, resulting in limited reference sequence databases that lack prior descriptions of Myxococcota-specific integrases.

Despite the strong co-occurrence of the *sul4* module with integron integrases, these elements are not mobile themselves (Gillings, 2017; Loot et al., 2024) and thus do not fully explain how *sul4* is translocated into new contexts and host bacteria. Instead, the ISCR element within the *sul4* module holds a prominent potential for this task (Peng et al., 2023; Razavi et al., 2017). In most *sul4* modules investigated here, the detected ISCR element represented one of the most recently described members of this group, ISCR28 (Yuan et al., 2024). Due to the *ori*IS and *ter*IS sequences and the rolling circle mechanism they enable (Toleman et al., 2006), these elements can translocate themselves and, in some cases, their adjacent sequences into new contexts (Yuan et al., 2024). Interestingly, Yuan and colleagues described a diverse range of genetic contexts for those ISCR28 elements where intact *ori*IS and *ter*IS sequences were present, while with an incomplete *ori*IS, the wider contexts of ISCR28 were highly conserved (Yuan et al., 2024). The authors deduced that in the latter cases, ISCR28 was not mobile but rather a passive element, co-transmitted through the activities of other MGEs (Yuan et al., 2024). With this background, the intact *ori*IS sites in all ISCR28 in our *sul4* modules together with the fact that they were found in a wide variety of genetic backgrounds and host bacteria would support its mobile nature in this context. Conversely, finding *attC* sites at or right after in most ISCR28 would imply that these elements are passively shuffled as components of the integron cassettes adjacent to *sul4* modules. If ISCR28 would be mobile itself, the question remains as to whether it also mobilizes adjacent genes, namely *sul4*. In our data, the arrangement of ISCR28 and *sul4* in one effluent read resembled the sandwich-like structure reported previously enabling the translocation of another ARG (Yuan et al., 2024). In this structure, a remnant ISCR28 with additional *ori*IS exceeds also upstream *sul4,* and the fusion by the two sequential *ori*IS leads to the capture of the sandwiched ARG, in our case *sul4* (Yuan et al., 2024).

Finding other than ISCR elements, namely members of various IS families, in three different *sul4* modules, suggests that a broader range of IS elements than previously thought could facilitate the transfer of *sul4* into new hosts, which is worrying given the substantial abundance and diversity of IS elements found in bacterial genomes (Kirsch et al., 2024). However, as all these three sequences with non-ISCR28 IS elements were found in *sul4* hosting plasmids, it may be that the IS elements are passively co-transmitted throughout the plasmid-mediated spread of *sul4*. Although we found evidence supporting the role of both integrase and ISCR28 for the *sul4* module mobilization, there is still no definite conclusion for this mechanism.

Besides the presence of integrases and ISCR28 flanking *sul4* and their absence in the same species MAGs lacking *sul4*, the most compelling evidence for the mobility of *sul4* module lies in its presence across various contexts, including plasmids, chromosomal regions, diverse genetic backgrounds, and even among unrelated species. Two of the newly discovered *sul4* hosts found in this study – *DUVQ01 sp012840965* and *Trichlorobacter sp933509905* – represented phyla Myxococcota and Desulfobacterota. These phyla were only recently established during the reclassification of former members of the diverse and non-monophyletic Deltaproteobacteria class, previously placed under the Proteobacteria phylum (Langwig et al., 2022; Waite et al., 2020). As the cultivation procedures of these species are not straightforward (Van Den Berg et al., 2015; Zou et al., 2024) the advancements in culture-independent omics methods have increased our understanding of species diversity in these groups, as shown by the high number of uncultivated bacteria (UBA) in both phyla (Kurashita et al., 2024; Langwig et al., 2022). Notably, in the NCBI database, the taxonomy for accession GCA_012840965.1, which is the reference genome of Myxococcota *DUVQ01 sp012840965* assigned by GTDB-Tk, is classified under *Oligoflexus* sp. of the order Oligoflexales. This nomenclature reflects the old and recently revised classification scheme for members of the class Deltaproteobacteria (Langwig et al., 2022; Waite et al., 2020).

Desulfobacterota and Myxococcota are widespread in microbial communities of soils and various aquatic environments, such as marine and freshwater habitats as well as wastewater (Langwig et al., 2022; Murphy et al., 2021) and they are known for their diverse metabolic abilities and involvement in various nutrient cycles (Langwig et al., 2022; Murphy et al., 2021; Sorokin et al., 2023; Waite et al., 2020) For instance, the unusual roles of *Trichlorobacter ammonificans* G1 in nitrate reduction processes were recently described within an activated sludge inoculum (Sorokin et al., 2023). In addition to the versatile secondary metabolite production (Kurashita et al., 2024), Myxococcota are most recognized for their active predatory members, whose functions within activated sludge communities in wastewater have been extensively studied (L. Zhang et al., 2023). In contrast to protist predators, Myxococcota are more prey-selective, favoring for instance *E. coli* and *Pseudomonas putida* (L. Zhang et al., 2023) and other gram-negative bacteria (Morgan et al., 2010). This could play a considerable role in the dissemination of resistance genes such as *sul4,* within a microbial community as the proximity between bacterial cells facilitates horizontal gene transfer (Cooper et al., 2013).

Similarly to Desulfobacterota and Myxococcota, the members of the Chloroflexota phylum, represented by the third *sul4* carrier bacterium found in our wastewater, are frequently encountered in diverse aquatic environments, including marine ecosystems, as well as in wastewater, particularly activated sludge (Björnsson et al., 2002; Murphy et al., 2021; Petriglieri et al., 2023). Diverse biochemical cycles have been described also among these bacteria (Narsing Rao et al., 2022). Intriguingly, based on the phylogenetic analysis of Sul4 and other similar DHP synthase protein sequences across bacterial lineages, in context of the first discovery of *sul4*, Razavi and colleagues proposed further investigations for studying Chloroflexota as the putative original host of *sul4* (Razavi et al., 2017) which supports our findings of Chloroflexota bacterium as one of the *sul4* carriers also in our samples.

None of the other *sul4* flanking regions examined in this study displayed similar genetic contexts to those found in *Trichlorobacter sp933509905* (Desulfobacterota), *DUVQ01 sp012840965* (Myxococcota) or *DSWT01* sp. (Chloroflexota) MAGs. Thus, the observation of phage sequences belonging to the ubiquitous (Du et al., 2023; Z. Li et al., 2021) ARG transmission associated (Wang et al., 2024) class Caudoviricetes near the *sul4* module in both, Myxococcota and Chloroflexota *sul4* MAGs raised the question about the phage’s putative role in mobilizing *sul4* across bacterial lineages. However, the comparison of two *DUVQ01 sp012840965* MAGs, with and without *sul4* module, revealed that a somewhat similar phage region was observed in both, suggesting that these sequences are unrelated to the presence and mobilization of *sul4*. It may also be that the region rich in phage sequences serves as a favorable target for the incorporation of other transconjugant genetic material (Dobrindt et al., 2004), such as via integrases and ISCR28. Likewise, the prophage sequence integration could occur at sites high in MGEs, explaining the similar structures observed in different genomes.

The *sul4-*encoding MAGs of all three species *DUVQ01 sp012840965* (Myxococcota)*, Trichlorobacter sp933509905* (Desulfobacterota), and *DSWT01* sp. (Chloroflexota) were found in samples from dried sludge. However, individual reads with *sul4* were also detected in the influent and effluent. These findings suggest that bacteria carrying *sul4* entered the wastewater treatment system via influent and potentially enriched in the activated sludge due to changes in environmental conditions modulating the bacterial composition and diversity (Dueholm et al., 2022; Ju et al., 2019). Moreover, the treatment conditions may have facilitated the transmission of *sul4* genes between different bacterial carriers. Finally, some of these genes or their carrier bacteria persisted throughout the treatment process as they were still detectable in the effluent and dried sludge. The elevated genus level bacterial diversity in the influent compared to dried sludge likely hindered the detection, or at least the metagenomic assembly of *sul4* contigs in the influent.

In addition to species of Myxococcota, Desulfobacterota, and Chloroflexota, other common wastewater species, such as *Sphingobium yanoikuyae* and *Desulfobacillus sp008933825*, were among the *sul4* hosts based on our analysis of sequences from public databases. Similarly to the PAH-degradation encoding genes found in the plasmid of *S. yanoikuyae* (Yin et al., 2020), the *sul4* gene within this plasmid could facilitate its rapid adaptation to new and diverse environmental stresses. Moreover, the putative aminoglycoside riboswitch identified within this *sul4* module refers to the regulation of ARG expression in response to aminoglycoside antibiotics and putative integration of further integron-encoded gene cassettes (J. Zhang et al., 2020). However, no additional ARGs were yet detected at this site.

Species within the *Desulfobacillus* genus, such as *D. denitrificans*, are known for their role in anammox consortia, where they contribute to nitrogen gas production through the anaerobic oxidation of ammonium in wastewater sludge (Nie et al., 2024). The adaptation mechanism of this consortia to antibiotic stress was recently described (Nie et al., 2024); Along with the subsequent recovery in nitrogen removal during long-term sulfamethoxazole exposure, a notable increase in the detection of *sul* genes was observed in time, and the greatest increase was seen for *sul4* (Nie et al., 2024). As the authors speculate, the increased production of modified DHP synthase encoded by the *sul* genes would unlock the folate pathway, inhibited by sulfamethoxazole (Nie et al., 2024). Moreover, the interspecies cooperation seemed to play a crucial role in this mechanism as evidenced by the increased frequency of MGE-mediated ARGs and the high genome plasticity enabling the transfer of adaptation-improving genes such as *sul* genes between the anammox consortia members and other sludge bacteria (Nie et al., 2024), which would further widen the host range of these genes in wastewater.

Conclusively, given the long half-life and low removal efficiency of sulfamethoxazole in traditional wastewater treatment processes (Kovalakova et al., 2020), the presence of antibiotic residues could provide favorable conditions for accumulation, spread across environmental and clinically relevant bacteria or even the emergence of novel *sul* variants. Depending on the intended use and discharge destination of effluent and dried sludge, the *sul4* genes and *sul4*-carrying bacteria found in them may spread further in the receiving microbial communities increasing the environmental burden of human-induced resistance. Moreover, high concentrations of sulfonamide antibiotic residues have been measured in natural waters especially in low- and middle-income countries (Ariyani et al., 2024; Razavi et al., 2017; Zhu et al., 2024) where the use of sulfonamides is pronounced (WHO, 2018) and the treatment of wastewater less regulated or efficient (UN Habitat and WHO, 2021). These conditions, resembling those in the original detection of the *sul4* gene in India (Razavi et al., 2017), could give rise to the further emergence and spread of *sul* variants among environmental bacteria and potentially species relevant to human health.

Besides taxa related to wastewater, we identified several other *sul4* carriers among opportunistic human pathogens. To our best knowledge, the patient isolate *S. enterica* subsp. *enterica* from China in 2020 is so far the only bacterial isolate for which the phenotypic resistance of *sul4* has been proven (Peng et al., 2023). This multidrug-resistant strain belonged to *S. enterica* serovar Thompson, a common cause of human infections and foodborne outbreaks (Zhao et al., 2021). Curiously, Peng and colleagues demonstrated that the *sul4* gene in this strain was integrated into the chromosome from a hybrid plasmid consisting of type IncHI2/HI2A common in gram-negative pathogens and gene arrays rich in ARGs and MGEs (Peng et al., 2023). The evidence of *sul4* circulating in plasmids of common pathogenic species is concerning as any new modes of dissemination for *sul4* will enhance the rapid spread of this gene. In addition to the prior plasmid phase of the *sul4* module in *S. enterica* and plasmid-encoded *sul4* in *S. yanoikuyae* (Yin et al., 2020), we identified two other distinct clinical *Aeromonas* spp. strains’ plasmids carrying *sul4*. Moreover, the additional ARGs within these *sul4-*containing integron cassettes could facilitate the co-selection resulting in the simultaneous mobilization of multiple ARGs (Nielsen et al., 2022). Since at least one metagenomic read from our influent wastewater dataset matched the genetic context of the *Aeromonas* plasmids, we speculate that this *sul4* carrier was present also in our samples, although we did not manage to assemble these reads into contigs or MAGs to support this conclusion.

Among the remaining wider contexts of *sul4*, we found matching genome sequences classified as *Moraxella osloensis* and *Leucothrix sargassi* strains (Liu et al., 2019) in both influent and effluent wastewater reads. However, some uncertainty remains about the taxonomic identity of these strains. The contamination in the latter reference genome suggested that in fact, both *sul4* hosts belonged to the *Moraxella* genus. Unfortunately, we were unable to assemble any contigs to construct MAGs from the matching wastewater reads in our dataset, to confirm the species identity of these *sul4*-carrying bacteria. However, despite the lack of taxonomical confirmation, finding these highly similar sequences in different data sets and geographical locations supports the widespread prevalence of this *sul4* module context.

In summary, we show that the recently recognized sulfonamide resistance gene, *sul4* is carried by diverse bacteria representing various ecological niches and taxonomical lineages. The highly conserved *sul4* module, containing *fol*K, *sul4,* and an ISCR28 was linked to different integron integrase genes in all investigated sequences, suggesting that the module is involved in the mobilization of *sul4*, although, the exact mechanism remains to be fully confirmed. Beyond these elements, the possibility of additional mobility-inducing mechanisms, such as the transfer of plasmids, co-selection through integron cassettes with multiple ARGs, or potential phage-mediated transmission raises further concerns regarding the effectiveness of crucially important sulfonamide antibiotics in human and animal medicine. Species involved in the essential biochemical cycles for nutrient removal in activated sludge, such as Myxococcota, Desulfobacterota, *Desulfobacillus*, and *Sphingobium* were found to be significant contributors to the carriage of the *sul4*. Therefore, the role of activated sludge in possibly enriching the host bacteria of sulfonamide resistance should be investigated in more detail. As many of these bacteria are also considered environmental taxa, our results provide further evidence for the ongoing discussion about the role of environmental bacteria in the currently seen emergence and spread of ARGs, for instance, occurring at wastewater treatment plants, where bacteria from different sources are mixed (Berglund et al., 2023; Finley et al., 2013; Forsberg et al., 2012). Moreover, this study demonstrates that the *sul4*-carrying bacteria or their genes can pass through wastewater treatment and end up in treatment outlets highlighting the need for ARG monitoring of WWTP discharges.

## Acknowledgments

This work was supported by the Research Council of Finland funding for the Multidisciplinary Center of Excellence in Antimicrobial Resistance Research. M.M. received funding from the MBDP doctoral program at the University of Helsinki. We would like to acknowledge CSC, IT Center for Science, Finland, for providing the computational resources for the study and the DNA Sequencing and Genomics Laboratory (supported by HiLIFE and Biocenter Finland funding), Institute of Biotechnology, University of Helsinki for sequencing and especially Pia Laine for the expertise in PacBio sequencing. Finally, we wish to thank Salla Pankka for her contribution to the preliminary analysis. Open access was funded by the Helsinki University Library.

List of Supplementary Materials:

## Supplementary Methods

### Supplementary Tables

Sheet 1: geNomad

Sheet 2: iPHoP

Sheet 3: Phage genes

Sheet 4: ISCR elements

### Supplementary Results

Supplementary Figure 1: Contigs included in methylation analysis

Supplementary Figure 2: Bacterial diversity by Metaphlan4

Supplementary Figure 3: Phylogenetic analysis of ISCR elements

